# Classical-contextual interactions in V1 may rely on dendritic computations

**DOI:** 10.1101/436956

**Authors:** Lei Jin, Bardia F. Behabadi, Monica P. Jadi, Chaithanya A. Ramachandra, Bartlett W. Mel

## Abstract

A signature feature of the neocortex is the dense network of horizontal connections (HCs) through which pyramidal neurons (PNs) exchange “contextual” information. In primary visual cortex (V1), HCs are thought to facilitate boundary detection, a crucial operation for object recognition, but how HCs modulate PN responses to boundary cues within their classical receptive fields (CRF) remains unknown. We began by “asking” natural images, through a structured data collection and ground truth labeling process, what function a V1 cell should use to compute boundary probability from aligned edge cues within and outside its CRF. The “answer” was an asymmetric 2-D sigmoidal function, whose nonlinear form provides the first normative account for the “multiplicative” center-flanker interactions previously reported in V1 neurons (Kapadia et al. 1995, 2000; Polat et al. 1998). Using a detailed compartmental model, we then show that this boundary-detecting classical-contextual interaction function can be computed with near perfect accuracy by NMDAR-dependent spatial synaptic interactions within PN dendrites – the site where classical and contextual inputs first converge in the cortex. In additional simulations, we show that local interneuron circuitry activated by HCs can powerfully leverage the nonlinear spatial computing capabilities of PN dendrites, providing the cortex with a highly flexible substrate for integration of classical and contextual information.

**Significance Statement:** In addition to the driver inputs that establish their classical receptive fields, cortical pyramidal neurons (PN) receive a much larger number of “contextual” inputs from other PNs through a dense plexus of horizontal connections (HCs). However by what mechanisms, and for what behavioral purposes, HC’s modulate PN responses remains unclear. We pursued these questions in the context of object boundary detection in visual cortex, by combining an analysis of natural boundary statistics with detailed modeling PNs and local circuits. We found that nonlinear synaptic interactions in PN dendrites are ideally suited to solve the boundary detection problem. We propose that PN dendrites provide the core computing substrate through which cortical neurons modulate each other’s responses depending on context.

## Introduction

In the primary visual cortex, the main feedforward pathway rises vertically from the input layer (L4) to the next stage of processing in layer 2/3 (Jennifer S. Lund, Angelucci, & Bressloff, 2003). These “driver” inputs establish the L2/3 pyramidal neurons (PNs) classical receptive fields, but account for only a small fraction of the excitatory contacts innervating those cells (<10%) (Binzegger, Douglas, & Martin, 2004). A much larger number of contacts (>60%) (Binzegger et al., 2004; Stepanyants, Martinez, Ferecskó, & Kisvárday, 2009) arises from the massive network of horizontal connections (HCs) through which cortical PNs exchange contextual information (Angelucci et al., 2002; Bosking, Zhang, Schofield, & Fitzpatrick, 1997; Boucsein, 2011; Chisum, Mooser, & Fitzpatrick, 2003; McGuire, Gilbert, Rivlin, & Wiesel, 1991; Rockland & Lund, 1982). Despite their large numbers and undoubted importance, relatively little is known regarding the HC’s contributions to behavior, the functional form(s) of the classical-contextual interactions they give rise to, or the biophysical mechanisms that underlie their modulatory effects.

Object boundary detection provides an attractive framework for studying classical-contextual interactions in visual cortex, given that object contours are known to contain essential information for recognition (Biederman, 1987; Field, Hayes, & Hess, 1993), and neurons in the first visual cortical area already show strong boundary-related contextual modulation effects(C.-C. Chen & Tyler, 2001; Chisum et al., 2003; Kapadia, Ito, Gilbert, & Westheimer, 1995; Kapadia, Westheimer, & Gilbert, 2000; Nelson & Frost, 1985; Polat, Mizobe, Pettet, Kasamatsu, & Norcia, 1998) (Figure 1a). With the aim to link this behaviorally relevant computation to underlying neural mechanisms, we followed a “normative” approach (Barlow, 1961; Laughlin, 1989; Olshausen & Field, 1996), founded on two mild assumptions: (1) that the goal of some V1 neurons is to detect object boundaries in natural scenes; and (2) that those neurons have access through their intracortical connections to certain boundary cues both within and outside their CRFs. Committing to these assumptions allowed us to interrogate natural images in a systematic way, and in so doing to determine what function a neuron should use to compute boundary probability from the available cues. This natural image-derived “classical-contextual interaction function” (CC-IF) served two purposes: (1) it helped put existing neurophysiological data on firmer theoretical ground, by providing “reasons” for nonlinear receptive field interactions that have been previously observed, and (2) it helped constrain the search for underlying neural mechanisms, by pointing to a specific function that V1 circuitry may need to compute, subject to the usual constraints of efficacy, biological plausibility, and parsimony.

**Figure 1.**
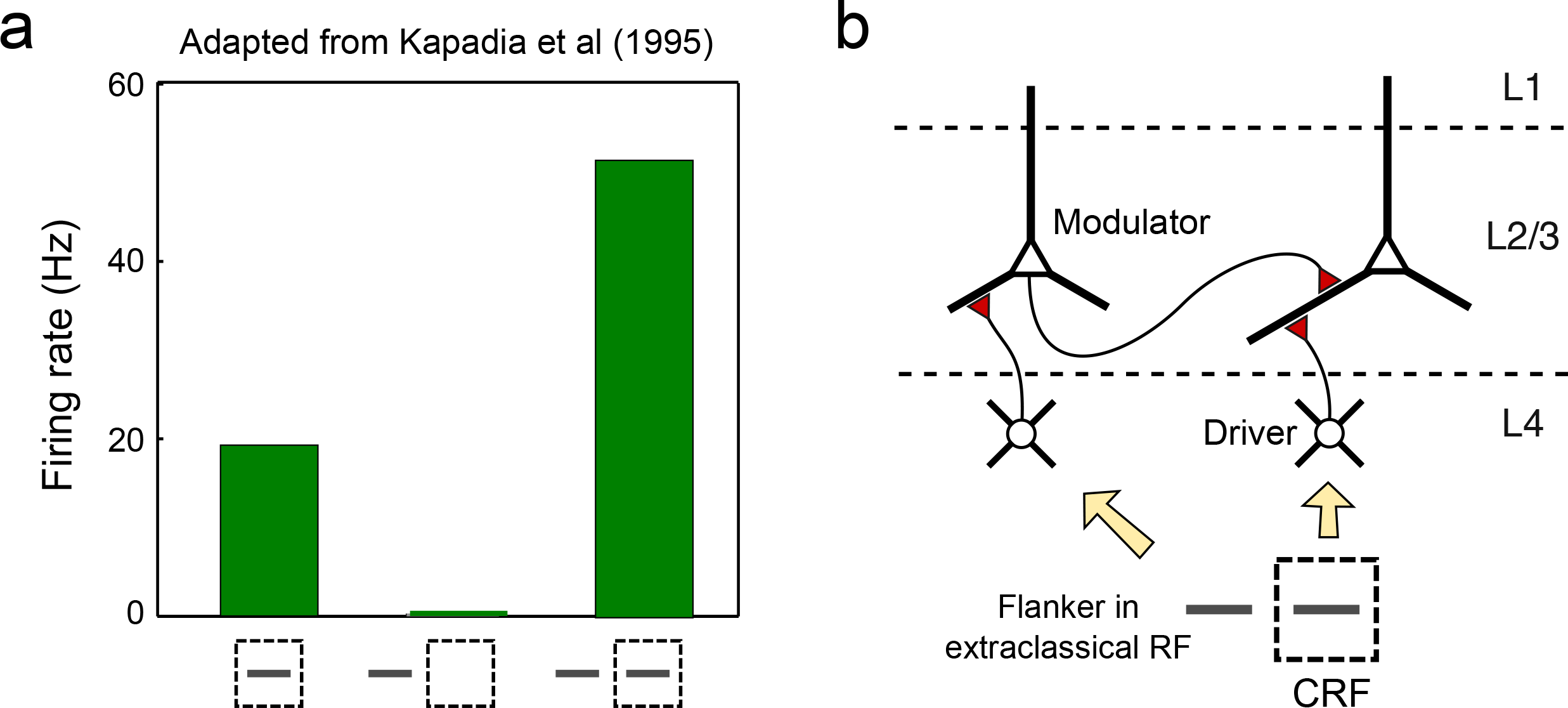
Classical contextual interactions in V1: phenomenon and proposed conceptual circuit. **a.** Schematic of key result of Kapadia et al. (1995). Cell recorded in monkey V1 showed modest response (~20Hz) to bar stimulus in CRF (dashed box); ~no response to flanking bar stimulus in surround, but strongly boosted response when flanking bar was paired with the CRF stimulus. **b**. Circuit model potentially accounting for the “multiplicative” classical-contextual interaction in **a**. Driver input representing the CRF stimulus rises vertically from layer 4, terminating with a distal bias on basal dendrite of a layer 2/3 PN. Horizontal input from neighboring V1 cell representing the flanker stimulus terminates on same PN dendrite with a proximal bias. From Behabadi et al (2012), the proximal modulator is expected to multiplicatively boost the dendrite’s response to the driver input.

The three main stages of the work were as follows. First, to characterize the computing problem faced by a putative boundary-detecting neuron in V1, we collected oriented filter responses on and off object boundaries in human-labeled natural images. From this data we constructed the first (that we know of) natural image-derived CC-IF that captures how the contrast levels of aligned boundary elements within and outside a neuron’s classical receptive field should be combined to determine object boundary probability. As described below, the CC-IF, which we found to have an asymmetric 2- D sigmoidal form, provides the first natural image-based explanation for three hallmark features of boundary related classical-contextual interactions previously reported in the neurophysiological literature (Kapadia et al., 1995, 2000; Polat et al. 1998).

Second, we noticed that the asymmetric 2-D sigmoidal form of the boundary-detecting CC-IF is qualitatively similar in form to the nonlinear 2-D input-output functions resulting from NMDAR-dependent spatial interactions between synapses targeting proximal vs. distal sites on a PN thin dendrite (Behabadi, Polsky, Jadi, Schiller, & Mel, 2012) (Figure 1b). We therefore attempted to fit the input-output behavior of a detailed neuron model to the natural image-derived CC-IF, as a test of the hypothesis that PN dendrites could be the neural substrate where boundary-related CC-IFs are computed in V1. We show that nearly perfect fits could be achieved, supporting the idea that nonlinear synaptic integration effects in PN dendrites could contribute to classical-contextual processing in V1.

Third, we carried out prospective simulations to assess the generality and expressive power of this dendrite-based spatial/analog computing mechanism. This was essential since the exact form of the boundary-related CC-IF we collected is tied to specific assumptions about (1) the inputs available to the pyramidal neuron – in our case, two aligned contour elements, one within, and the other outside the cell’s CRF), and (2) the task the neuron is supposed to perform based on those inputs – which in our case was calculating boundary probability. Since either of these assumptions could have been different, leading to a different CC-IF, it is critical to identify sources of flexibility in the cortex that could allow HCs to produce a wide variety of classical-contextual interactions. To this end we used compartmental models to explore the spectrum of CC interactions achievable through variations in single neuron and circuit-level parameters. We conclude that PN dendrites, forming the core of the cortical circuit, provide a powerful computing substrate through which HCs can flexibly modulate neural responses depending on context.

## Material and Methods

### Natural image labeling

The pair-difference filter was directly inherited from an earlier study (Zhou & Mel, 2008) with all parameters unchanged. Image patches were selected based on their PD filter response centered at regularly spaced values {0, 0.1, 0.2…1.0} along each of the two filter dimensions, with a bin width +/− 0.005 Image patches were collected until each of the 121 bins contained at least 30, but no more than 100 patches. The image patches were displayed in a 21-inch monitor when shown to the labeling participants through a MATLAB program. The patches were shown with a red box, representing the center receptive field (CRF) and the center dot as shown in Figure 2.2b. The labeler was told to follow the following rules to evaluate the existence of a contour within the center receptive field (given as printed instructions to them):

1. an object contour was present in the red box.
2. the contour entered one end of the box and exited the other while always remaining within the box.
3. the contour was unoccluded at the box center, indicated by the red dot.

**Figure 2.**
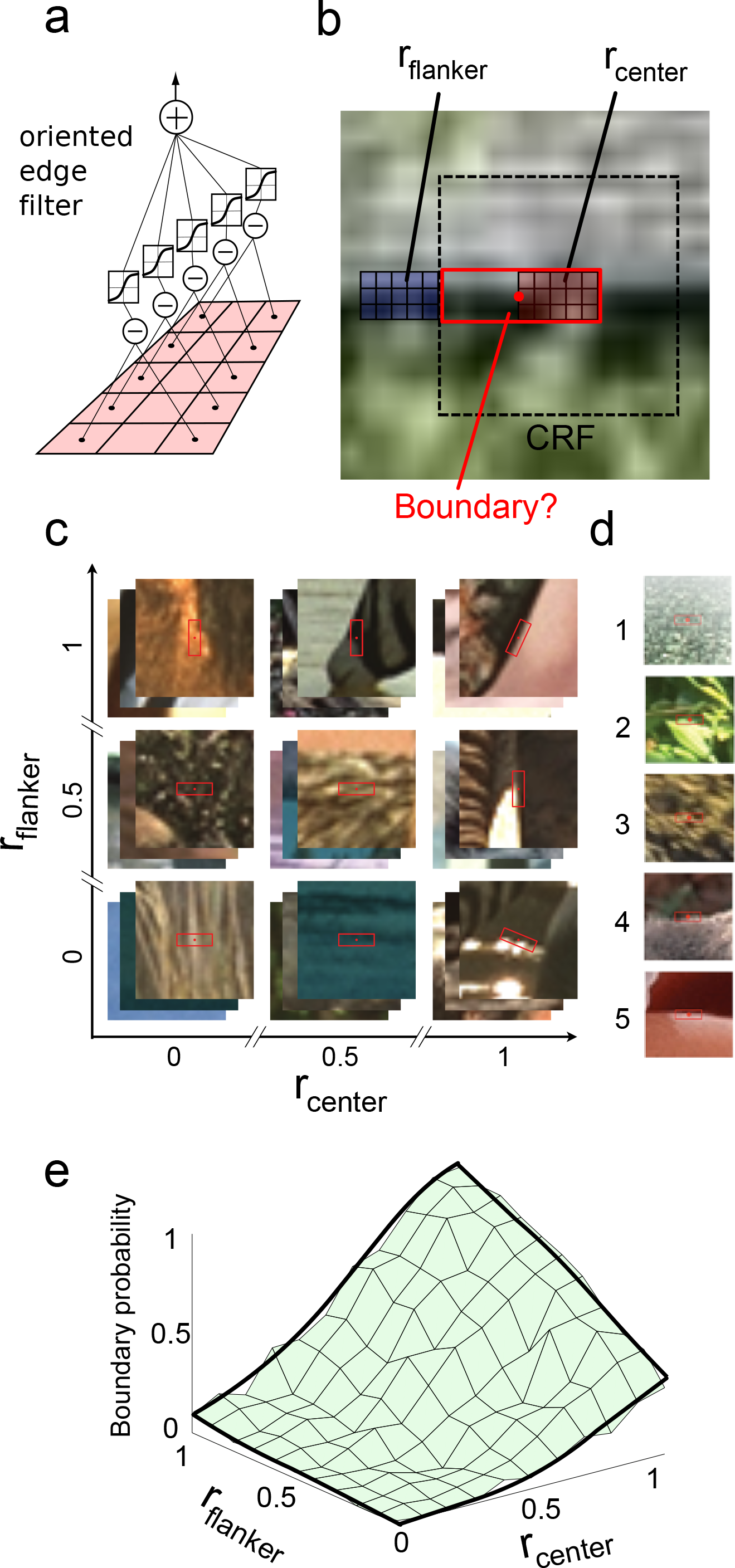
Measurement of a classical-contextual interaction function (CC-IF) from human-labeled natural images. **a.** Schematic of oriented edge filter. The filter response was obtained by computing pairwise differences PDs for five pixel pairs, each separated by an unsampled pixel. Each PD was passed through a sigmoidal nonlinearity, given by x/(0.2+x) for x≧0, and x/(0.2-x) for x<0, and the results were summed. Sign indicated edge polarity. **b.** Sample image patch shown with the CRF of a virtual V1 neuron superimposed (dashed box). Each patch was characterized by responses of two aligned filters r_center_ within the CRF, and r_flanker_ just outside the CRF. **c.** Image patches were drawn at random from a natural image database and binned based on r_center_ and r_flanker_ values, forming a 2-dimensional space of bins. Only 3 of the 11 bins are shown along each axis. **d.** Human labelers were asked to judge whether an object contour was present in the red box that entered one end, exited the other, remained always within the box, and was unoccluded at the center. Scores were assigned as follows: 1 = “definitely no”, 2 = “probably no”, 3 = “can’t decide”, 4 = “probably yes”, 5 = “definitely yes”. Examples of patches that received each label are shown. **e.** Boundary probability within each image bin was plotted as a function of r_center_ and r_flanker_.

The labeler also received instructions to give scores in the following way:

1. No contour
2. Not likely a contour - some structured elements were seem within the CRF but not likely forming a contour going through it
3. Likely a contour - contour seen but either occluded at the center of the CRF, or is not aligned with the orientation of the CRF
4. Almost certainly a contour - contour seen but occluded at non-center positions or is slightly curvy.
5. Surely a contour with aligned orientation

The labeler pressed 1-5 on the keyboard to score each patch and the data were recorded automatically. The patches were shown in pseudorandom order. If the labeler made a mistake they could stop the program by pressing esc and get back to the last patch they labeled to change their score. Each experimental session lasted for about an hour, during which the labeler was able to label 600-1,000 different patches. A total of 16,000 labels were collected to generate figure 2.2c.

### Compartmental simulation

Simulation were run within the NEURON simulation environment (version 7.5 standard distribution). Unless otherwise specified, the compartmental model, biophysical parameters and ion channel parameters regarding the NMDAR AMPAR and GABA-A were the same as in two earlier studies ((Behabadi et al., 2012) Table 1 and (M. Jadi et al., 2012) Table 2). A 3D-reconstructed layer 3 pyramidal neuron morphology ((Amatrudo et al., 2012), cell name “Jul16IR2b-V1”, source from Neuromorpho.org) was used in producing the fitting results in figure 3. For the rest of the paper results, a layer 5 PN morphology from prior studies (“j4”) was used (Behabadi et al., 2012), which made it easy to compare to the prior results. A Gaussian current was injected at the soma with a mean current of 1.0 nA and standard deviation of 0.75 nA. Under this condition the output was linearly correlated with the current flowing to the soma in range of 0-150 Hz. This was a proven-efficient way to remove the somatic nonlinearity. Neuron files are available upon request.

**Figure 3.**
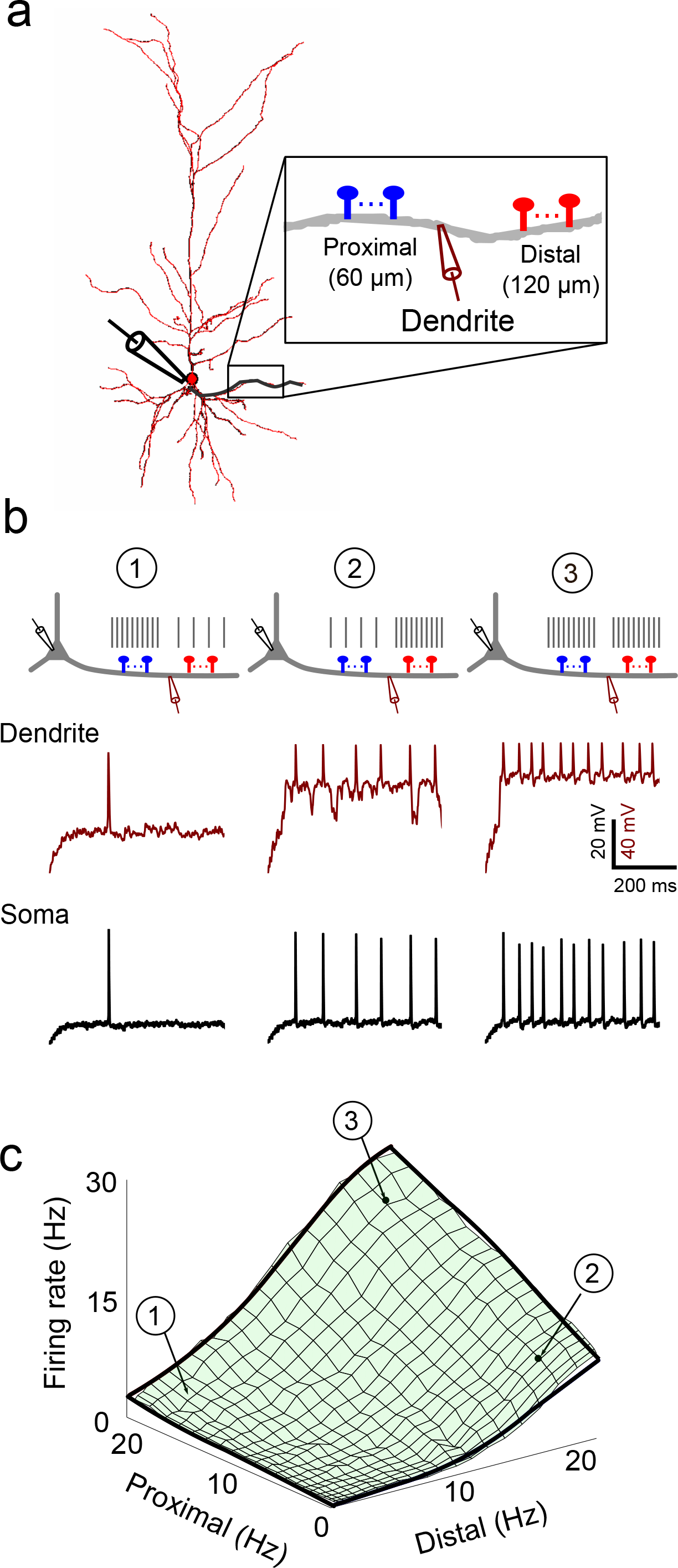
Responses of a biophysically-detailed compartmental model of a layer 2-3 PN in response to two spatially-separated excitatory inputs delivered to a basal dendrite. Model neuron was from macaque V1 (see methods for detail). **a.** Two excitatory input “pathways”, each consisting of 30 identical excitatory spine synapses, were placed in clusters at 60 and 120 μm from the soma. Each spine contained both an NMDA and an AMPA-type conductance (see Methods), and was stimulated by a regular spike train (with random phase) with frequency ranging from 0-20 Hz. Voltage was recorded at the soma and in the stimulated dendrite 90 μm from the soma. **b.** Dendritic and somatic recordings for 3 representative cases: (1) 17.5 Hz proximal and 2.5 Hz distal simulation. (2) 2.5 Hz proximal and 17.5 Hz distal; and (3) both proximal and distal at 17.5 Hz. All traces started from −70 mV. **c.** Firing rates were averaged over 10 1-second runs. Bold outer frame is identical to that in Figure 2e.

In generating figure 3c, the x and y axis were elongated according to the relationship x’ = x^1.5^. The motivation was to model synaptic depression effects, in which the effect of the stimulus input with linearly increasing presynaptic frequency may only climb up sub-linearly.

Spines were modeled as two cylindrical compartments. Unless otherwise specified, the morphology was as follows: neck, cylinder shape, with a height of 1 μm and a diameter of 0.05 μm; head, cylinder shape, with a height of 0.5 μm and a diameter of 0.5 μm. The two compartments were then attached to a parent dendrite. In figure 2.5c, when simulating the “increased neck resistance” case, the diameter of the neck compartment is set to 0.025μm. Only excitatory synapses (AMPAR/NMDAR) were modeled with spine morphology. Inhibitory synapses (GABA-A) were modeled directly innervating the dendritic shaft without spines.

**Figure 4.**
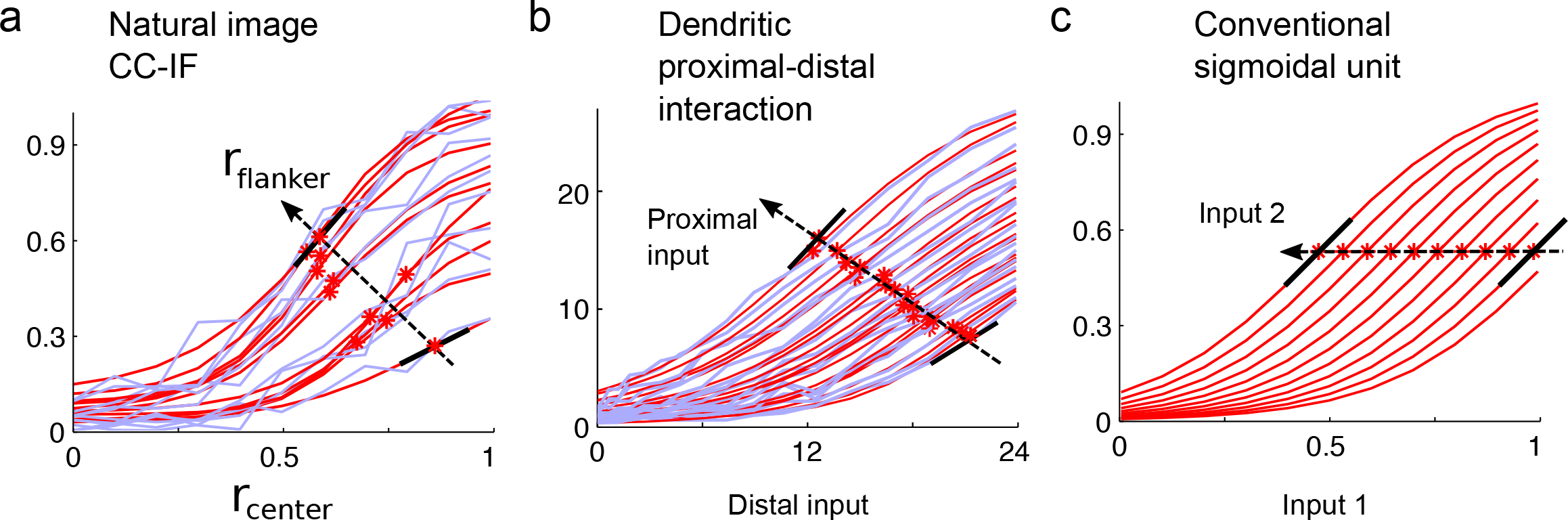
Nonlinear structure of the image-derived CC-IF compared to a dendritic proximal-distal interaction function, and a conventional sigmoidal neural activation function. **a.** Each CC-IF slice (blue) was fit by a logistic function (red) with variable threshold, slope, amplitude, and y-offset. Steepest slope of each fit is marked by an asterisk; black bars at lowest and highest modulation levels help visualize progression of threshold and slope values. Progression of inflection points has a significant non-zero slope (two sided test, p<0.01) **b.** Same as **a** but for responses of compartmental model with a distal driver and proximal modulatory input. Slices are plotted over a larger range of inputs than in **a** for better visualization of the sigmoidal form. **c**. Progression of i/o curves of a conventional sigmoidal activation function with two inputs; peak slopes remain constant regardless of “modulation” level.

**Figure 5.**
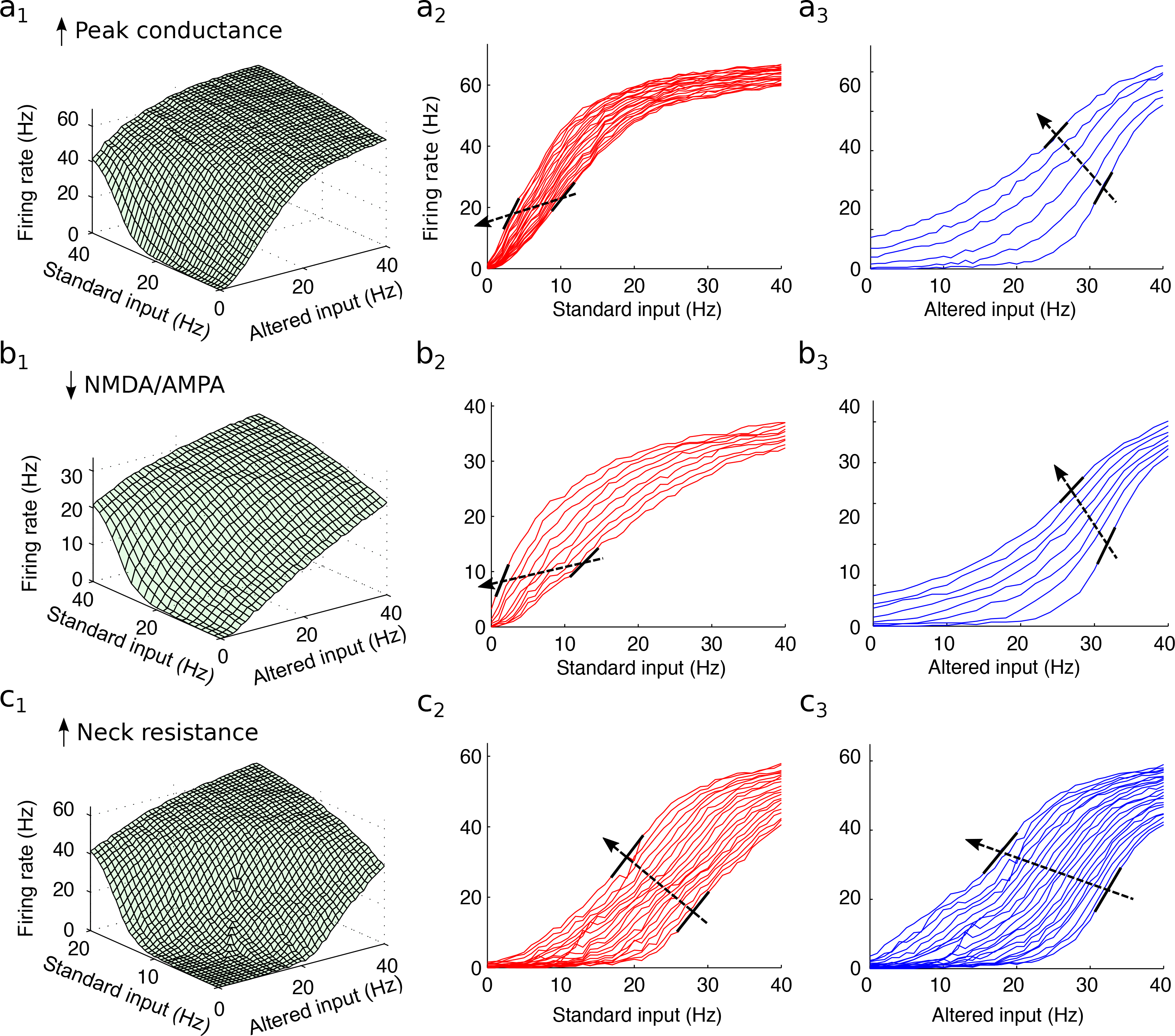
Analysis of dendritic response asymmetries due to differences other than spatial bias. In all cases, both Standard and Altered inputs were located 60 μm from the soma. Only one synaptic parameter was changed in the Altered input; Standard input was always the same. **a**. Peak synaptic conductance was increased 3-fold in the Altered input. Panel a_1_ shows the 2-D response surface; panels a_2_ and a_3_ show slice views of the same data. Black bars again show steepest points of individual i/o curves. Dashed arrow shows direction of increasing modulatory level. **b**. Same as **a** but Altered input has NMDA/AMPA ratio lowered from 2 to 0.1 Total conductance was increased to produce roughly the same amount of activation compared to the Standard input. **c.** Same as a but Altered input had spine neck resistance increased 3.5 fold to ~500MΩ.

All simulations were done with computational resources of the University of Southern California high performance computing center.

## Results

### Deriving a contour-related CC-IF from human labeled natural images

In studying the boundary-related responses of V1 neurons, a typical observation is that a cell’s response to an oriented contour element inside its classical receptive field (CRF) is boosted (often 2-3 fold) by aligned “flankers” lying outside the CRF, whereas the flankers produce little or no response on their own (C. C. Chen, Kasamatsu, Polat, & Norcia, 2001; Kapadia et al., 1995, 2000; Polat et al., 1998)(Figure 1a). This nonlinear facilitatory interaction between center and flanker stimuli concords well with psychophysical effects (Kapadia et al., 2000; Polat & Sagi, 1994), but also seems intuitive, in that evidence for an object boundary within a cell’s CRF ought to be “amplified” when corroborated by evidence from nearby locations. Our progress in understanding the biophysical mechanisms underlying this type of classical-contextual interaction has been hampered, though, by the lack of a method for quantitatively predicting the form of the CC-IF under different assumptions about a neuron’s goal and available inputs. The ability to predict CC-IFs would be valuable in two ways: it would provide a reference to which a measured CC-IF could be compared, and a target towards which biophysical modeling efforts could be aimed.

With this goal in mind, we turned to natural images to obtain an empirical CC-IF involving two aligned boundary elements, one inside and one outside the CRF of a virtual V1 neuron (by analogy with the stimulus configuration of Kapadia et al. 1995 - see Figure 1a). We first constructed a 3×5 pixel oriented edge filter (Figure 2a) loosely inspired by the receptive field structure of a canonical even-symmetric V1 “simple cell” (Hubel & Wiesel, 1962). The filter returns a value r ∈ [0,1] when applied at any position/orientation in an image, signifying the strength of the oriented luminance contrast at that site. Image patches (100×100 pixels) were collected at random from the Corel image database, and a CRF (dashed rectangle in Figure 2b), was placed (virtually) at the center of each image patch (i.e., was not actually drawn). Two filter values were computed for each patch: r_center_, measured within the virtual CRF, and r_flanker_ from an aligned position just outside the CRF (Figure 2b). Image patches were then sorted based on this pair of measured filter responses. An 11×11 grid of image bins was defined over the 2-D space of center-flanker response pairs. These bins were centered at regularly spaced values {0, 0.1, 0.2…1.0} along each of the two filter dimensions, with a bin width +/−0.005. Image patches were collected until each of the 121 bins contained a minimum of 30, but typically 100 image patches. A few of the image bins are illustrated schematically in Figure 2c. Image patches that did not fall into any bin were discarded. To collect responses from human labelers, patches in each of the 121 bins were presented on a video monitor in pseudorandom order. Labelers were told to focus on a red box contained within the CRF (Figure 2b), and asked to assign a score ranging from 1 to 5 (without time pressure) indicating their level of confidence that all of the following were true: (1) an object contour was present in the red box; (2) the contour entered one end of the box and exited the other while always remaining within the box; and (3) the contour was unoccluded at the box center, indicated by the red dot in Figure 2d. Scores were linearly converted to a [0,1] range and averaged within each bin, yielding a plot of ground truth “boundary probability” over the 2-D space of center-flanker score pairs (Figure 2e).

The plot in Figure 2e illustrates three features typical of a CC-IF(Kapadia et al., 1995). First, as the value of r_center_ increases when r_flanker_ is near zero, boundary probability rises to a modest level (40.5%). In contrast, as the value of r_flanker_ increases when r_center_ is near zero, boundary probability remains low (8.9%). Despite this weak effect on its own, a strong flanker input more than doubles the gain of the response to the center input, leading to a boundary probability of 98.3% when both center and flanker filters are strongly activated.

While other spatial configurations of two or more filters, or different filter designs, or different labeling criteria could have been used, it is notable that an arrangement of just two aligned filters used to sort image patches into bins, coupled with a simple definition of an object boundary, already allowed us to capture the hallmark features of a contour-related CC-IF.(Kapadia et al., 1995; Polat et al., 1998)

### Could PN dendrites be the site where the CC-IF is computed?

What biophysical mechanisms might be capable of producing this peculiar type of functional interaction? The fact that horizontal and vertical inputs first converge on the basal and apical oblique dendrites of layer 2/3 PNs raises the question as to what role PN dendrites might play in nonlinearly integrating classical and contextual inputs. A previous combined neurophysiological and modeling study (Behabadi et al., 2012) showed that NMDAR-dependent interactions between proximal and distal synapses on PN basal dendrites produce asymmetric 2-D sigmoidal interaction functions similar to the plot of Figure 2e (see also M. P. Jadi, Behabadi, Poleg-Polsky, Schiller, & Mel, 2014). This similarity led us to ask whether a spatially-biased projection pattern in which horizontal axons project closer to the soma where they can exert a more multiplicative effect, and vertical inputs from layer 4 connect more distally (Figure 1b), could reproduce the boundary-related CC-IF we had extracted from natural images (Figure 2e).

To test this, we ran compartmental simulations of a PN stimulated by groups of 30 proximal and distal synapses (Figure 3a). Examples of simultaneous dendritic and somatic recordings are shown in Figure 3b for 3 different input intensity combinations. Dendritic traces show either no active response (case 1), intermittent slow dendritic spikes (case 2), or plateaus (case 3), with fast back-propagating somatic action potentials superimposed. We systematically varied proximal and distal input rates from 0 to 20 Hz, and recorded the firing rate at the soma. The resulting 2-D neural response function (Figure 3c) closely matched the natural image-derived boundary probability plot shown in Figure 2e; the bold outer frames are identical in the two plots. All three hallmark features of a classical-contextual interaction were again present: the distal input alone drove the cell to fire at a moderate rate (11.02 Hz). In contrast, the proximal input alone drove the cell to fire only weakly (3.16 Hz), but significantly boosted the gain of the response to the distal input when the two pathways were activated together (25.96 Hz at full activation). Thus, we concluded that the analog nonlinear processing capabilities of PN dendrites are well suited to produce the asymmetric interaction between CRF and extra-classical inputs for purposes of boundary detection in natural images.

### On the nature and potential sources of the neural response asymmetry

Could the asymmetry of the CC-IF be produced by a different (and especially a simpler) mechanism, not involving dendritic spatial integration? A common abstraction of a neuron (or dendrite’s) input-output function is a weighted sum of inputs followed by a sigmoidal nonlinearity,

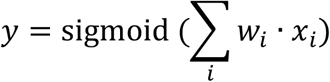

This conventional neural activation function produces a nonlinear interaction between its inputs *x*_*i*_ by virtue of the output nonlinearity, and can even produce an asymmetric nonlinear interaction by assigning different weights (*w*_*i*_) to each input. Can such a model, properly parameterized, capture the type of asymmetric nonlinear interaction expressed by the natural image-derived CC-IF, eliminating the need to consider more complex dendritic integration mechanisms? To address this question, we fit each iso-modulator slice of the CC-IF with logistic functions whose threshold, slope, and asymptote were allowed to vary arbitrarily (see Figure 4 caption for formula and parameters used). As shown in Figure 4a, the best-fitting sigmoids (red curves) followed a progression wherein, as the threshold (i.e. x-coordinate of the steepest point) moved leftward under the influence of an increasing modulatory input, both the maximum slope and amplitude of the sigmoidal curves increased. These correlated changes in threshold, slope, and amplitude are summarized graphically by the upward-leftward shift and steepening of the black bars marking the maximum slopes moving from low to high modulation levels. The same progressive increase in slope and amplitude was seen in the proximal-distal dendritic interaction function (Figure 4b; note the i/o curves are plotted over a greater range of inputs than in Figure 4a to more fully visualize the curves’ sigmoidal form). The existence of changes in slope and amplitude from curve to curve rules out that the CC-IF can be represented by a conventional sigmoidal activation function, for which the maximum slope and amplitude of the i/o curves remain unchanged across modulation levels (Figure 4c; this is true regardless of which input is considered the driver and which the modulator). Based on these results, we conclude that the natural image-derived CC-IF has nonlinear structure that falls outside the representational scope of a conventional neural activation function.

Having established that a conventional sigmoidal activation function lacks the fundamental asymmetric structure needed to represent the natural image-derived CC-IF, we next asked whether a proximal-distal separation of driver and modulator pathways on PN dendrites is *necessary* to achieve the type of asymmetric nonlinear interaction seen in the CC-IF, or whether other types of asymmetries between the two input pathways can produce a similar type of interaction.

While it would be impossible to consider all possible alternatives, we identified three types of differences between two pathways, other than dendritic location, that could potentially produce the type of amplitude+slope boosting interaction seen in the CC-IF. We then ran simulations in which: (1) the classical and contextual synapses were co-mingled on the dendrite, thus eliminating their spatial asymmetry, (2) one of the pathways retained its original biophysical characteristics and was called the “Standard input”; and (3) the other pathway was altered in one of three ways:

1. <u>Increased peak synaptic conductance.</u> *Rationale*: Increasing the peak conductance of an input pathway effectively lowers its threshold for NMDAR activation, which could lead to an enhanced superlinear interaction between the two pathways;
2. <u>Reduced NMDA-AMPA ratio.</u> *Rationale*: by making one (e.g. the driver) pathway less superlinear on its own, it might benefit more from the nonlinear excitability boost provided by the modulatory pathway;
3. <u>Increased spine neck resistance.</u> *Rationale*: higher spine neck resistances amplify spine voltages, and are said to “encourage electrical interaction among coactive inputs” and “promote nonlinear dendritic processing” (Harnett, Makara, Spruston, Kath, & Magee, 2012).

The results of these manipulations are shown in Figure 5. In each row, the surface plot at left shows the 2-D interaction with the “Standard” input (same in all 3 rows) plotted on the left abscissa and the “Altered” input plotted on the right abscissa. The 2-D interaction surface is shown sliced along both cardinal directions in the middle and right plots of each row, and the maximum slope points of the slices are again marked by black bars. While the interaction functions take on various forms, in none of the cases or for either direction of slicing do we observe conjoint amplitude+slope increases with increasing value of the modulator. Based on these negative results, we conclude that the peculiar type of nonlinear interaction between two inputs that arises from a dendritic location asymmetry, which closely matches the nonlinear structure of the CC-IF, cannot be easily reproduced by simply modifying the relative excitability of one of the two pathways in the absence of a dendritic location asymmetry.

### Interneuron circuits provide additional flexibility for tailoring classical-contextual interactions

Importantly, the natural image-derived CC-IF (Figure 2e) that has guided our search for underlying mechanisms is just one of an essentially unlimited number of different interaction functions that could be needed in different cortical areas, which must process very different kinds of information, and in different animal species, which must perform well in very different kinds of environments. We therefore set out to more fully explore the spectrum of CC-IFs that could be produced by varying anatomical and physiological parameters available to cortical neurons and circuits. As a first step we generated dendritic interaction functions that deviated in various ways from the standard shown in Figure 3c (reproduced in Figure 6a over for larger range of inputs), achieved by: (1) reducing the separation distance of the classical and contextual inputs from 90 to 30μm (Figure 6b); (2) altering the NMDA conductance model to eliminate post-synaptic receptor saturation (Figure 6c); and (3) increasing spine neck resistance from 100 MΩ to 500 MΩ. In all three cases, unlike those in Figure 4, synapses in both the driver and modulator pathways were identical but for their dendritic location. The gallery of cases in Figure 6 illustrates that even when driver and modulator synapses have identical properties, different spatial biases in their projections to PN thin dendrites can produce CC-IF’s that are widely varying in functional form.

**Figure 6.**
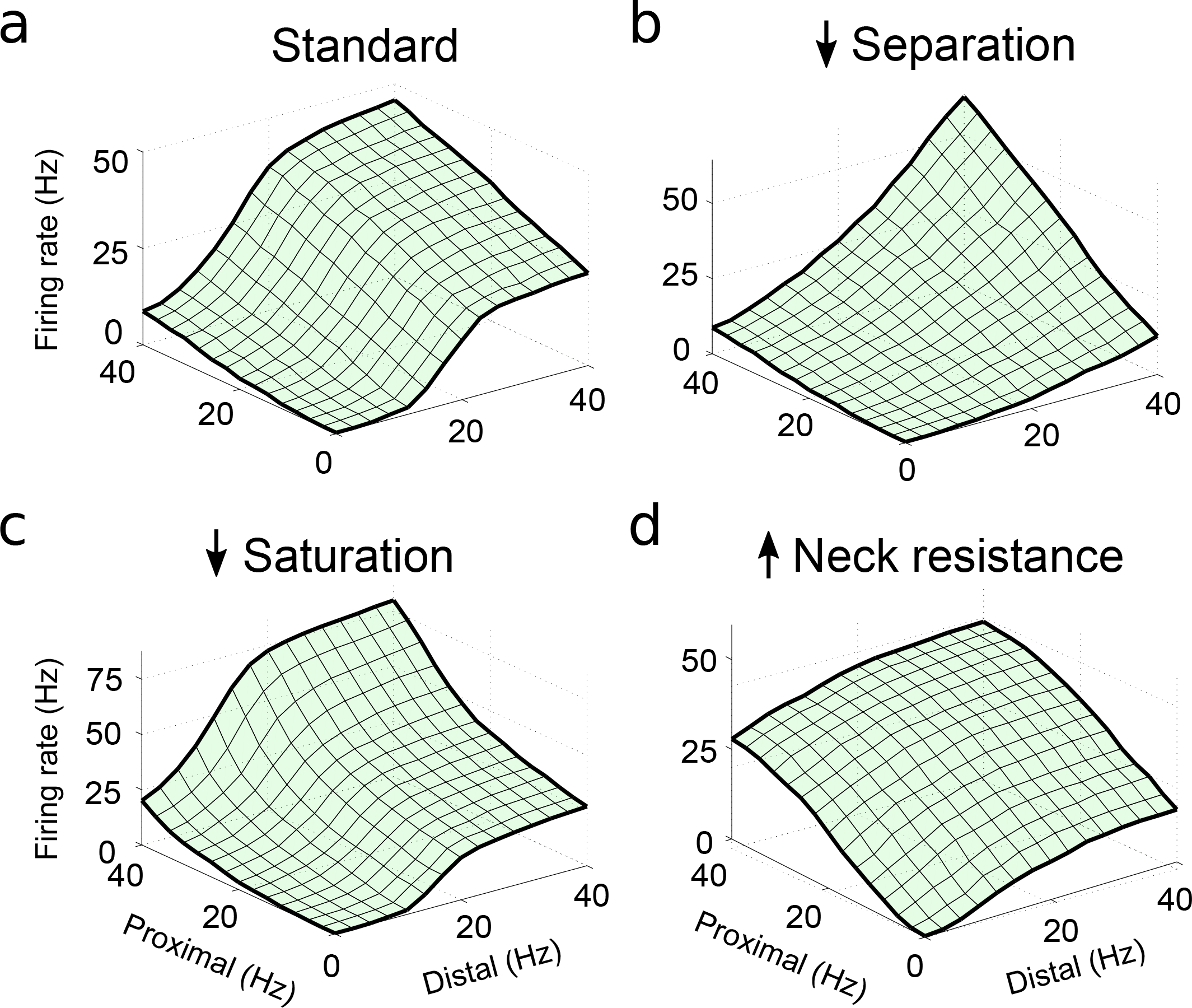
Variations in dendritic response functions resulting from changes in parameters of the compartmental model. **a.** 2-D response surface for a standard seed condition (proximal-distal separation = 90 μm, spine neck resistance = 100 MΩ, synaptic peak conductance = 2 nS, and synaptic saturation “cap” was 100% (meaning that a single presynaptic release event saturated all available channels at the synapse; a cap of 200% meant that through repeated stimulation, temporal facilitation could produce up to twice the base conductance; a cap of ∞ meant synapse did not saturate at any rate. **b.** Spatial separation of the two inputs was reduced to 30 μm. **c.** Synaptic saturation cap was set to ∞. **d.** Spine neck resistance was set to ~500 MΩ.

An additional source of flexibility that could be used by the cortex to tailor classical-contextual interactions lies in the parameters that govern how inhibitory interneurons activated by horizontal axons affect PNs both directly and indirectly. We focused on the circuit motif shown in Figure 7a, a subset of the full interneuron circuit summarized in Pfeffer, Xue, He, Huang, & Scanziani (2013) and Tremblay, Lee, & Rudy (2016). Our reason for focusing on the SOM->PV->PN subcircuit stems from the fact that SOM interneurons are strongly activated by HCs (Adesnik, Bruns, Taniguchi, Huang, & Scanziani, 2012), and are therefore particularly relevant for understanding contextual modulation. Furthermore, SOM interneurons have the interesting property that they inhibit PN dendrites directly, but also inhibit PV interneurons, which leads to a *disinhibition* of PNs perisomatically. The net effect of this arrangement is that activation of HCs produce a shift of inhibition away from the soma and towards the dendrites of their target PNs (Pfeffer et al., 2013). What might be the functional role of this local PV-SOM circuit motif?

**Figure 7.**
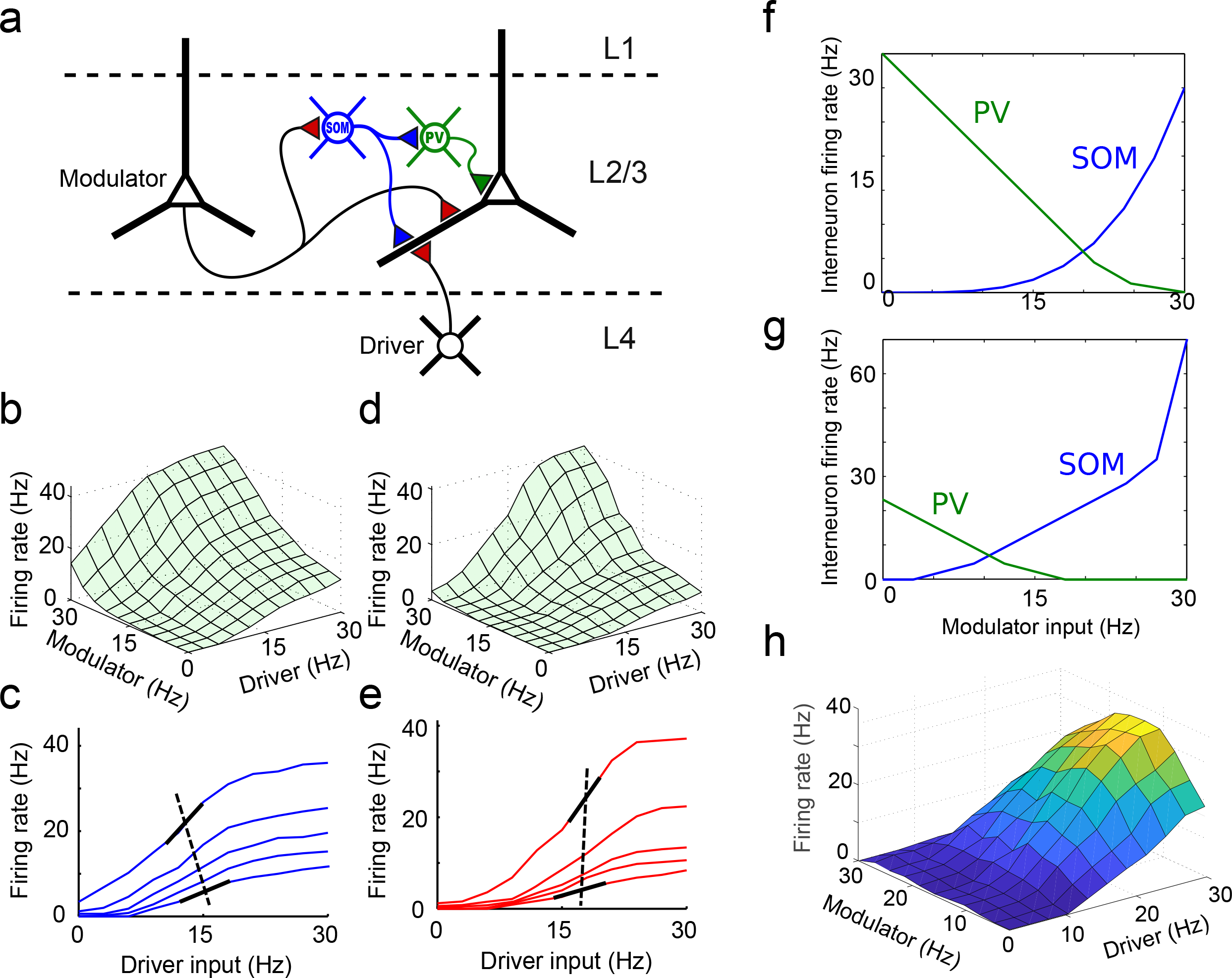
Dendritic spatial processing capabilities can be leveraged by local inhibitory circuits. **a**. Conceptual model modified from Figure 1b to include a SOM->PV interneuron circuit. SOM interneuron inhibits PN dendrite distally; PV interneuron inhibits PN perisomatic region, and is inhibited by SOM interneuron. SOM neuron is driven by horizontally offset modulator. **b.** Dendritic response function with pure excitatory spatial interaction, similar to Figure 3c with input range extended. **c.** Slices of surface in **b** (only even numbered lines are shown.) **d, e.** Same as **b** and **c** but with inhibitory effects included. Progression of i/o curves follows a relatively pure multiplicative scaling (as evidenced by aligned thresholds) with increased dynamic range. **f.** Activation curves of PV (green) and SOM (blue) interneurons used for the simulation shown in **d** and **e**. **g, h.** An alternative set of PV and SOM activation curves, providing an example of a non-monotonic modulatory effect.

Previous work has shown that excitatory (E) and inhibitory (I) synapses interact in complex ways in dendrites, depending on their absolute and relative locations.(Gidon & Segev, 2012; M. Jadi, Polsky, Schiller, & Mel, 2012; Koch, Poggio, & Torre, 1983; Vu & Krasne, 1992) Extrapolating from the findings of (M. Jadi et al., 2012), who focused on proximal-distal E-I interactions in active dendrites, we expected that increasing SOM activation by HCs should have two specific effects on a PN’s dendritic input-output curves, namely: (1) a progressive increase in the threshold (i.e. right-shifting) of the cell’s dendritic i/o curves, caused by the increasing distal inhibition; and (2) a progressive increase in the effective gain of the cell’s i/o curves, caused by the cell’s gradual alleviation from proximal (i.e. perisomatic) inhibition. As a “control” condition, the 2-D response surface produced by the compartmental model with no inhibition is shown in Figure 7b, and the corresponding 1-D slices are shown in Figure 7c (consisting of a subset of those in Figure 4b). The response surface with intact SOM+PV inhibition is shown in Figure 7d, and the corresponding 1-D slices are shown in Figures 7e.

Interestingly, when a modulatory input engages the interneuron subcircuit, leading to progressive threshold and gain increases in the dendritic input-output curves (as shown in Figure 7d,e), the net effect can be to produce (1) a more purely multiplicative scaling of the dendritic response curves, as evidenced by the more vertical alignment of the thresholds in Figure 7e compared to Figure 7c, along with (2) an expanded dynamic range from low to high modulation levels, as evidenced by the greater vertical spread of asymptotes in the red vs. blue slices. The functions that map the modulation intensity to PV (perisomatic) and SOM (dendritic) inhibitory firing rates in this example are shown in Figure 7f, and a detailed analysis of the separate and combined effects of the SOM and PV inhibitory components can be found in Figure S1. It is important to note that the two curves shown in Figure 7f were designed through trial and error to achieve a pure multiplicative effect, so that pure multiplicative modulation is by no means an inevitable outcome of this type of circuit, nor are the curves in Figure 7f likely to be unique in producing a multiplicative effect. Rather, the example is intended to illustrate the flexibility that even a simple interneuron circuit adds to an already richly expressive classical-contextual modulation capability based on pure excitatory dendritic location effects (Figure 4). A final example shows that modulation of a PN’s responses by a horizontal pathway is not restricted even to monotonic (i.e. facilitating or suppressive) effects: the SOM and PV activation curves shown in Figure 7g, modified relative to those in Figure 7f, lead to the non-monotonic 2-D interaction surface shown in Figure 7h. This type of interaction function might be appropriate in cases where (1) a driver input provides evidence for a particular preferred feature within the cell’s CRF (call it feature A), and (2) a contextual input provides contextual support for A up to a point, which increases the cell’s responsiveness to its primary driver input; but after that point, begins to provide stronger contextual support for a (non-preferred) feature B, which eventually reduces the cell’s responsiveness to its primary driver input.

To summarize, we have shown that the boundary-related CC-IF derived from natural images has a functional form which is outside the representational scope of a conventional neural activation function, but is readily produced by NMDAR-dependent spatial interactions in PN dendrites - at the very site where classical and contextual signals first converge in the cortex. Probing this mechanism in greater depth, we showed that variations in multiple synapse-related parameters, including spine neck resistance, peak synaptic conductance, degree of synaptic saturation, degree of synaptic facilitation, and NMDA-AMPA ratio, greatly expands the space of CC-IFs that can be produced by PN dendrites based on purely excitatory synaptic interactions. Finally, we showed that the spectrum of realizable classical-contextual interaction functions is further enriched when local interneuron circuits are taken into consideration. As examples, we showed activation curves for SOM and PV interneurons that lead to pure multiplicative modulation, and others that produce more complex, non-monotonic forms of modulation (Figure 7).

## Discussion

Our overarching goal in this work has been to gain insight into how the massive network of horizontal connections in the cortex provides a means for PNs to modulate each other’s responsivity to their primary CRF driver inputs through the back and forth exchange of contextual information. To gain traction, we focused on the problem of boundary detection, and on the mechanisms that V1 neurons may use to combine classical and contextual cues for this purpose. Why boundary detection, and why V1? Detecting object boundaries is a core process in biological vision, and is a crucial precursor to rapid object recognition (Biederman, 1987; Potter, 1976). As for V1, psychophysical and neurophysiological studies suggest V1 is heavily involved in the early stages of boundary processing (Adini, Sagi, & Tsodyks, 1997; Angelucci et al., 2002; C. C. Chen et al., 2001; Dresp, 1993; Field et al., 1993; Grosof, Shapley, & Hawken, 1993; Grosof et al., 1993; Ito & Gilbert, 1999; Kapadia et al., 1995, 2000; Levitt & Lund, 2002; W. Li & Gilbert, 2002; Mizobe, Polat, Pettet, & Kasamatsu, 2001; Polat & Sagi, 1994; Sceniak, Ringach, Hawken, & Shapley, 1999) and the network of horizontal axons, by linking cells with similar orientation preferences (Bosking et al., 1997; Gilbert & Wiesel, 1989) and co-linear or co-circular receptive fields (Chisum et al., 2003; Schmidt, Goebel, Löwel, & Singer, 1997) seems “designed” with long-range contour integration in mind.

By way of motivating our approach, it is useful to consider what could and could not be learned from a conventional study of classical-contextual interactions in V1, such as the seminal study of Kapadia et al. (1995). In one of their key neurophysiological findings (whose results are schematized in Figure 1a), the authors showed that a ~40% of V1 neurons exhibit an asymmetric nonlinear “facilitatory” interaction between a stimulus in the CRF that acts as a driver and an aligned flanker in the extra-classical RF that acts as a modulator (i.e. does not drive the cell by itself, but “multiplies” the cell’s response to its driver input). What their study could not answer were two questions that correspond to our main findings here:

1. “What *should* be the interaction between the center and the flanker inputs?”
2. “What biophysical mechanisms are capable of generating the appropriate classical-contextual interaction?”

We discuss our efforts to answer each of these questions below.

### On the various uses of the natural image-derived CC-IF

Our approach to answering question 1 flows from the fact that, if we are willing to assume a neuron’s goal is to detect object boundaries, then through a mechanical ground-truth labeling process applied to natural images, we can determine precisely what function the neuron *should* use to detect boundaries based on a given set of cues. The result of this process for the center and flanker cues shown in Figure 2b is the CC-IF shown in Figure 2e.

Having the natural image-derived CC-IF in hand provides two major benefits. First, the CC-IF creates a solid link between a classical-contextual interaction of a type that has been reported in the neurophysiological literature (Kapadia et al., 1995) and a specific natural sensory classification problem that the function helps to solve. This is the first normative account for a classical-contextual interaction in the cortex that we are aware of. Second, in relation to question (2) above, the CC-IF provides a well-founded target towards which biophysical modeling efforts can be aimed; this is how the CC-IF was used in the initial fitting exercise of Figure 3. Furthermore, the natural image-derived CC-IF contains sufficient detail and structure that it can help distinguish among competing mechanistic models; this is how the CC-IF was used in the analyses of Figures 4 and 5.

The CC-IF can be looked at in a third way: as a crude “algorithm” for detecting object boundaries, since it effectively scores every location in an image for boundary probability based on the responses of two aligned oriented filters. We would not expect the algorithm to perform well, given that it receives input from just two of a large number of filters in the neighborhood that could provide information about the presence or absence of an object boundary. (In contrast, full blown models of contour detection in V1 typically include inputs from many filters (Z. Li, 1998; Loxley & Bettencourt, 2011; Pettet, McKee, & Grzywacz, 1998; Ursino & La Cara, 2004; Yen & Finkel, 1998)). Nonetheless, when applied to natural images as a nonlinear filter in its own right, the CC-IF should at least show *some* capability for boundary detection. To verify this, we processed images with the local edge filter shown in Figure 2a, collected r_center_ and r_flanker_ values at each image location at 8 orientations and in the two complementary configurations shown in Figure 8a (in red and blue). We then used the two center-flanker value pairs as inputs to the CC-IF, and the scores obtained were averaged to yield a composite boundary probability measure at each location/orientation. Boundary images were generated by plotting the maximum boundary probability across all 8 orientations at each pixel. Examples of original, local edge, and boundary images are shown in Figure 8b-d. Higher probability is indicated by darker pixels. In comparison to local edge images, boundary images emphasized longer, well-formed object contours while suppressing textures, resulting in images that more closely resemble line drawings. These images in effect represent how neurons that compute the CC-IF would “see” the world.

**Figure 8.**
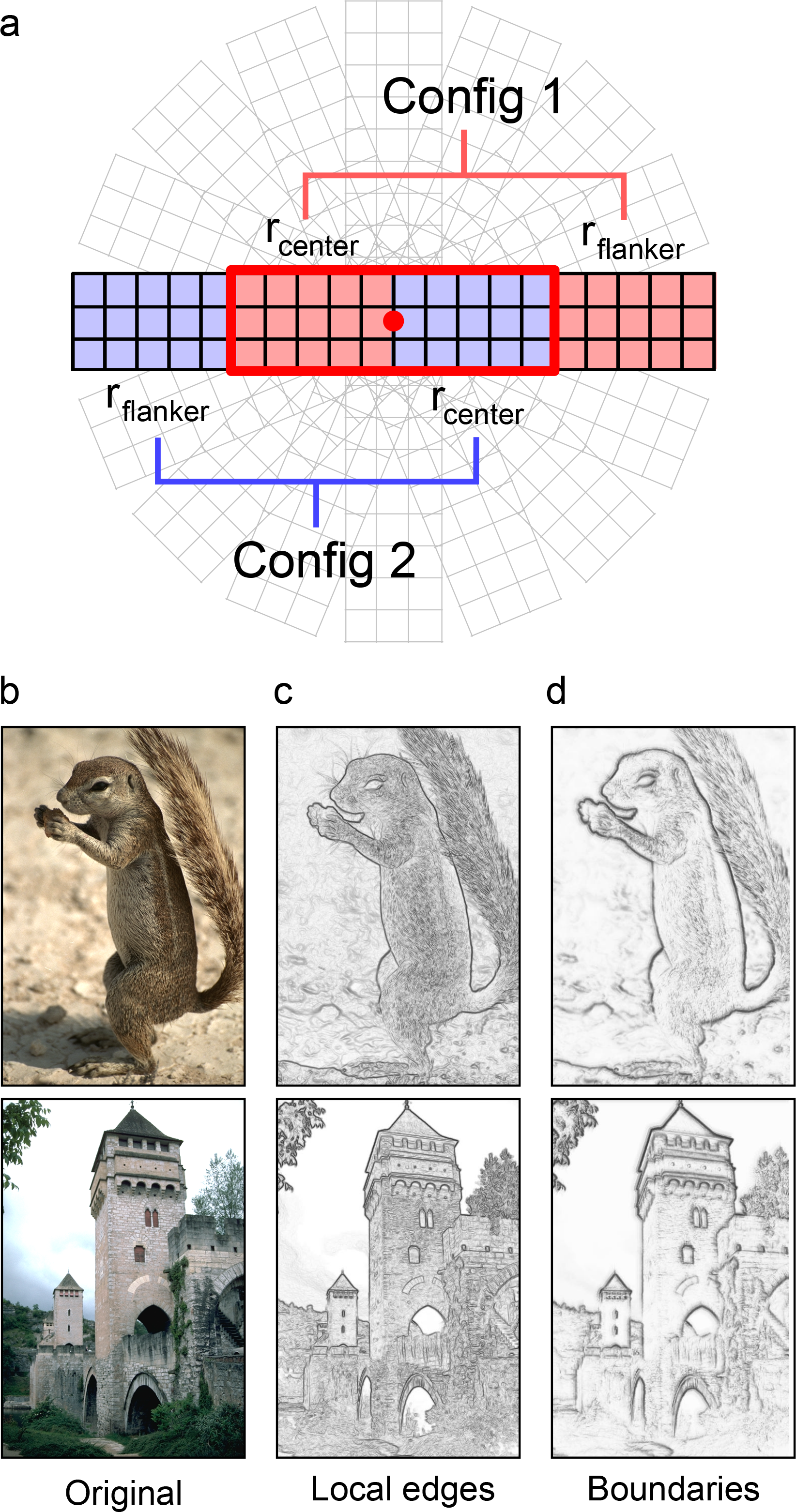
Applying the CC-IF as a nonlinear image filter. **a.** Two pairs (red and blue) of center and flanker edge filters in configurations like that shown in Figure 2b were evaluated at every pixel at 8 orientations (illustrated here for the horizontal orientation). Each pair of filter values was run through the CC-IF, and the results from the two configurations were averaged at each pixel/orientation. The contour response at each pixel was the maximum across all 8 orientations. **b.** Original images. **c.** Local edge images. Maximum edge score across all orientations at each pixel is indicated by the darkness of the pixel. **d.** Contour image. As for local edges, darker pixels indicated stronger contours. Contour images emphasized spatially-extended object contours and suppressed textures.

### Relationship to previous studies of object boundary statistics

Ours is not the first analysis of natural image statistics pertaining to object boundaries: Geisler et al. (2001) collected co-occurrence statistics of local boundary elements in natural images, and showed that they predict human contour grouping performance. While the image statistics collected by Geisler et al. also relate to object boundaries, the type of data they collected and their uses of that data are different. First, they began with a different premise: they assumed local boundary elements had already been detected. They then collected statistics such as (1) the probability that a second boundary element is found at all possible offsets in position and orientation relative to a first boundary element, and (2) the log likelihood ratio comparing the probability that a second boundary element at a given offset in position and orientation is part of the same or different object as a first boundary element. In contrast, our focus has been on the problem of discriminating object boundaries from non-boundaries at a given location based on a particular configuration of cues. This difference in objective is reflected in the different ways the natural image data is represented: our data is represented by a function, the CC-IF, which describes how two scalar oriented contrast measurements should be used to compute boundary probability at a location. The grouping statistics collected by Geisler et al. are represented as scalar values linking pairs of locations/orientations. In terms of its application, Geisler et al. used their data to explain human psychophysical phenomena, whereas we have used our data to help constrain neural models. In a study similar to that of Geisler et al., Sigman et al. (2001) also histogrammed the co-occurrence probabilities of pre-detected boundary elements at all position/orientation offsets from a reference boundary location. Their results were again expressed in terms of scalar values relating pairs of offset boundary elements. While interesting, their main conclusion – that boundary elements in natural images tend to lie on common circles – is not directly informative as to the computation needed to detect boundaries in the first place, nor to the neural mechanisms that may carry out those computations.

### Relationship to previous V1-inspired models of contour detection

Several V1-inspired models of contour detection have appeared in the literature (Z. Li, 1998; Loxley & Bettencourt, 2011; Pettet et al., 1998; Ursino & La Cara, 2004; Yen & Finkel, 1998), with one of two objectives (or both): (1) to explain human contour detection performance as a function of various stimulus parameters (e.g., element spacing, contour length, open vs. closed contours, density of distractors, etc.) (Z. Li, 1998; Pettet et al., 1998; Yen & Finkel, 1998), or (2) to perform well at detecting contours in noisy artificial and natural images (Z. Li, 1998; Loxley & Bettencourt, 2011; Ursino & La Cara, 2004). In all cases, these models were assembled using “off the shelf” neurally-inspired components and operations, including weighted sums, sigmoids, thresholds, divisive normalization and winner-take-all operations, etc. The core elements and operations that find their way into such models can generally be traced back to the earliest days of neural modeling (Rosenblatt, 1962; Rumelhart, Hinton, & McClelland, 1986), when a high premium was placed on the use of simple mathematical (or logical) operations. This long-established tradition explains the continued widespread use of weighted sums to represent synaptic summation, as well as the strong preference for nonlinearities produced by compact algebraic expressions.

In contrast to previous cortically-inspired models of boundary detection, the core computing operation of our model (represented by the CC-IF) was not assumed to take on any particular form, let alone a commonly used or mathematically compact one, but rather was derived from natural image data under the normative assumptions that object boundary detection was the task, and certain specific measurements were available to the neuron to solve the task. Interestingly, one of our key findings is that that core operation represented by the CC-IF is <u>not</u> readily expressible in the conventional neural vernacular, or with compact mathematical expressions of any kind. This challenge motivated our second main activity, which was to search for neural mechanisms capable of producing the unconventional type of nonlinear interaction we uncovered. Thus, proceeding from normative assumptions actively pushed us away from the simple mathematical operations used in most previous cortical models.

### Dendrites provide a parsimonious neural implementation of the CC-IF

One of our key findings is the close match between the image-derived CC-IF and the neural response function arising from NMDAR dependent proximal-distal synaptic interactions in PN dendrites. This match supports the prediction that PN basal and apical oblique dendrites contribute to contextual processing in V1, and perhaps elsewhere in the cortex, by providing a flexible analog computing substrate in which behaviorally-relevant nonlinear interactions between horizontal, vertical and potentially other input pathways can take place.

This prediction has three main preconditions: (1) appropriate physiological machinery; (2) appropriate anatomical connectivity; and (3) sufficient flexibility to support the tremendous range of computing capabilities that the cortex is evidently capable of providing (i.e. including multiple types of sensory processing, motor control, language, planning, emotion, etc.). Regarding the appropriate physiological machinery, a previous study carried out in brain slices established that NMDAR-dependent proximal-distal interactions in PN dendrites are *capable* of providing the type of asymmetric pathway interaction needed to fit the image-derived CC-IF (Behabadi et al., 2012; reviewed in M. P. Jadi et al., 2014). Regarding the appropriate anatomical connectivity, the first requirement is that classical (vertical) and contextual (horizontal) axons should target at least some of the same dendrites of PNs. Existing data strongly supports this: both horizontal and vertical axons terminate on PN dendrites throughout cortical layer 2-3 (Binzegger et al., 2004; Chisum & Fitzpatrick, 2004; Jennifer S. Lund et al., 2003; Yoshimura, Sato, Imamura, & Watanabe, 2000).The more demanding requirement is that horizontal and vertical axons project to PN dendrites with different spatial biases, especially along the proximal-distal extents of individual basal or apical oblique dendrites. Very little connectivity data is currently available at this sub-dendrite scale, though several observations, taken together, suggest that within-dendrite biases of this kind are biologically feasible: (1) the axonal projections of inhibitory neurons show famously strong spatial biases at the sub-dendrite scale (Bloss et al., 2016; DeFelipe, Ballesteros-Yáñez, Inda, & Muñoz, 2006; Karube, Kubota, & Kawaguchi, 2004; Tremblay et al., 2016); (2) excitatory pathways have well known spatial biases of other kinds, for example, they can selectively target dendrites in specific layers or parts of layers (Harris & Shepherd, 2015; J. S. Lund, 1988; Petreanu, Mao, Sternson, & Svoboda, 2009); (3) excitatory axons are subject to activity-dependent clustering, producing a tendency for co-activated axons to form contacts on nearby spines (DeBello et al., 2014; Iacaruso, Gasler, & Hofer, 2017; Lee, Soares, Thivierge, & Béïque, 2016; van Bommel & Mikhaylova, 2016, 2016; Weber et al., 2016); (4) individual excitatory axons can show strongly biased projections at the sub-dendrite scale (Bloss et al., 2018; Morgan, Berger, Wetzel, & Lichtman, 2016) (5) proximal vs. distal synapses can be subject to different plasticity rules, which could lead to a spatial sorting-out of functionally distinct input pathways (Froemke, Poo, & Dan, 2005; Gordon, Gribble, Syrett, & Granato, 2012; Sandler, Shulman, & Schiller, 2016); and (6) differences in EPSP rise times suggest horizontal vs. vertical axons (Yoshimura et al., 2000) and near vs. far horizontal connections onto pyramidal neurons (Schnepel, Kumar, Zohar, Aertsen, & Boucsein, 2014) do, on average, terminate at different distances from the soma.

Regarding the flexibility to produce a rich spectrum of CC-IFs in the cortex, as our compartmental simulations show, on top of the inherent spatial processing capabilities of PN dendrites, allowing variations in multiple synapse-related parameters, including spine neck resistance, peak synaptic conductance, degree of synaptic saturation, degree of synaptic facilitation, and NMDA-AMPA ratio, significantly expands the space of CC-IFs that can be produced by PN dendrites – even when limited to purely excitatory synaptic interactions. The spectrum of realizable CC-IFs is then greatly expanded when the parameters of local interneuron circuits are brought into play.

To summarize, PN dendrites have the appropriate capabilities, are located at the appropriate place, and are part of a circuit with the appropriate flexibility to contribute centrally to the integration of classical and contextual signals in V1, and potentially other cortical areas. Whether this powerful and flexible computing resource is used in the cortex for contextual processing remains an open question, but one that is answerable with currently available experimental techniques.

## Author contribution

L.J. participated in image labeling, compartmental modeling, data analysis, image processing, and writing of the manuscript. B.F.B. and M.P.J developed the compartmental simulation approach and contributed to the design of the study. C.A.R. developed the image labeling approach and contributed to the design of the study. B.W.M. contributed to the design of the study, and participated in data analysis and writing of the manuscript.

---

**Figure S1.**
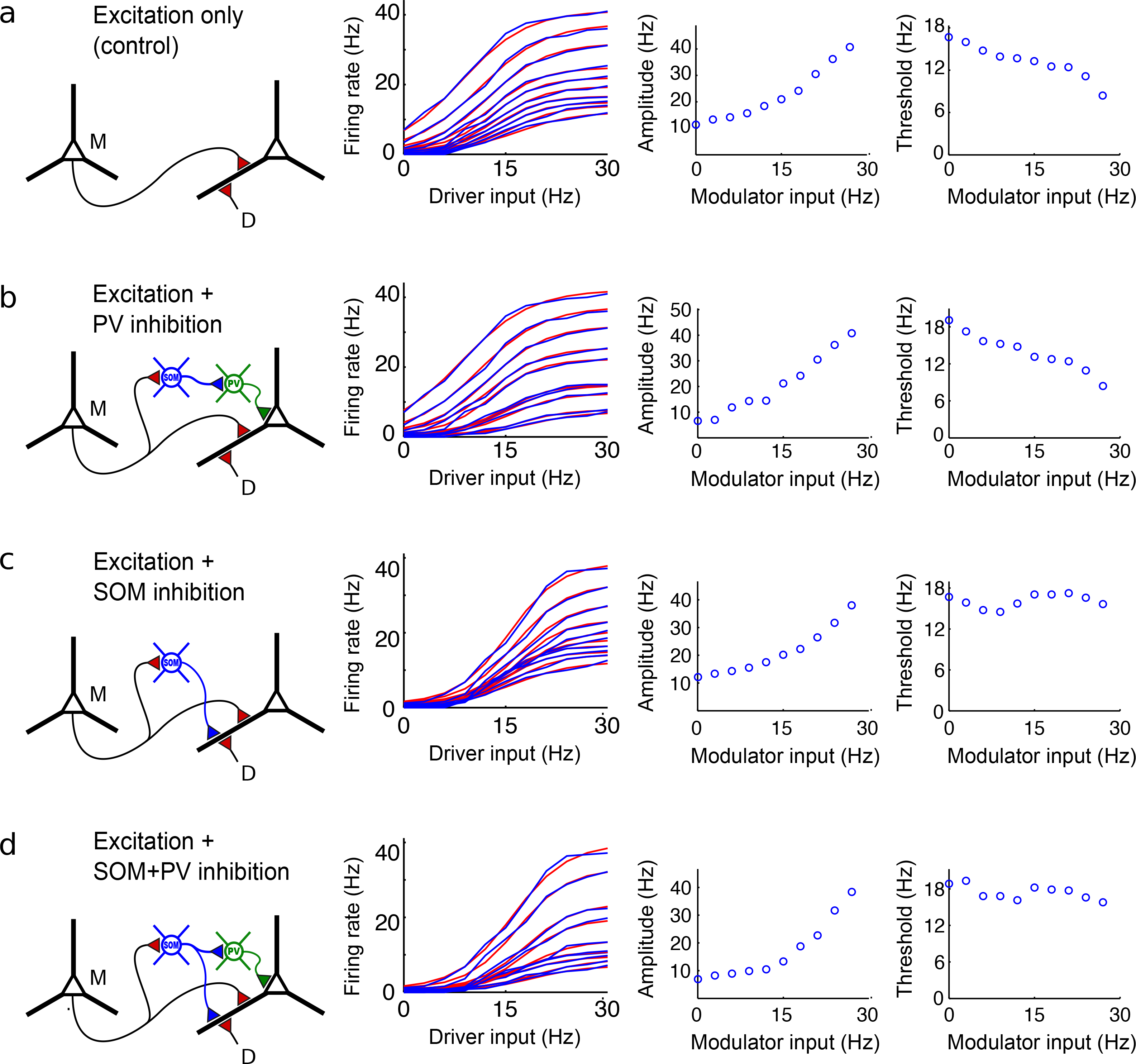
Elaboration of simulation run of Figure 7b-e leading to multiplicative scaling of i/o curves through combined effects of SOM and PV inhibition (and disinhibition). Each row shows circuit modeled; slices of 2-D response surface; and progressions of amplitude and threshold of best-fitting logistic function across modulation levels. **a.** Excitation only circuit, same as figure 7b. Amplitudes increase and thresholds decrease for increasing modulation levels. **b**. Circuit with disinhibition caused by inhibition of PV neuron by SOM neuron. Main effect is to increase dynamic range of response amplitudes as PV interneuron is progressively shut down. **c**. Circuit now including only dendritic inhibition by SOM interneuron. Main effect is threshold increase (and therefore gradual alignment of thresholds) across modulatory levels. **d.** Circuit combining both the dynamic range increase from b and threshold alignment from **c**. Effect is now a relatively pure multiplicative scaling of i/o curves over a large dynamic range, as in Figure 7**d-e**.

## Acknowledgements

This work was supported by the NIH/NEI BRP grant EY016093 and Israel-US BSF grant #2009341. The simulations were performed at the Center for High-Performance Computing of the University of Southern California.

